# Unveiling the Microbial Diversity and Associated Secondary Metabolism on Black Apples

**DOI:** 10.1101/2023.11.02.565319

**Authors:** Michael S. Cowled, Christopher B. W. Phippen, Kresten J. K. Kromphardt, Sidsel E. Clemmensen, Rasmus J. N. Frandsen, Jens C. Frisvad, Thomas O. Larsen

**Author notes:** Address correspondence to Thomas O. Larsen. Christopher B. W. Phippen, FMC Corporation, Genvej 2, DK-2970 Hørsholm, Denmark. Kresten J. K. Kromphardt, Unibio A/S, Asnæsvej 2A, DK-4400 Kalundborg, Denmark.

## Abstract

Black apples are the late-stage microbial decomposition of apples after having fallen to the ground. This phenomenon is highly comparable from year to year, with the filamentous fungus *Monilinia fructigena* most commonly being the first invader, followed by *Penicillium expansum*. Motivated by the fact that only little chemistry has been reported from apple microbiomes, we set out to investigate the chemical diversity and potential ecological roles of secondary metabolites (SMs) in a total of 38 black apples. Metabolomics analyses were conducted on either whole apples or small excisions of fungal biomass derived from black apples. Annotation of fungal SMs in black apple extracts was aided by cultivation of 15 recently isolated fungal strains on 9 different substrates in an OSMAC approach, leading to identification of 3319 unique chemical features. Only 6.8% were attributable to known compounds based on analysis of HPLC-HRMS/MS data using spectral library matching tools. Of the 1606 features detected in the black apple extracts, 32% could be assigned as fungal-derived, due to their presence in the OSMAC-based training dataset. Notably, the detection of several antifungal compounds clearly indicates the importance of such compounds for invasion of and control of other microbial competitors on apples. In conclusion, the diversity and abundance of microbial SMs on black apples was found to be much higher than that typically observed for other environmental microbiomes. Detection of SMs known to be produced by the six fungal species tested also highlights a succession of fungal growth following the initial invader *M. fructigena*.

**Importance:** Microbial secondary metabolites constitute a significant reservoir of biologically potent and clinically valuable chemical scaffolds. However, their usefulness is hampered by rapidly developing resistance, resulting in reduced profitability of such research endeavours. Hence, it is vital that the ecological role of such microbial secondary metabolites be considered to understand how best to utilise such compounds as chemotherapeutics. Here, we explore an under-investigated environmental microbiome in the case of black apples; a veritable “low-hanging fruit”, with relatively high abundances and diversity of microbially produced secondary metabolites. Using both a targeted and untargeted metabolomics approach, the interplay between metabolites, other microbes and the apple host itself was investigated. This study highlights the surprisingly low incidence of known secondary metabolites in such a system, highlighting the need to study the functionality of secondary metabolites in microbial interactions and complex microbiomes.

## Introduction

Black apples represent an unconventional ecological system: dynamic and never quite reaching a state of homeostasis, but following a development path that reoccurs year-on-year.(1, 2) These apples harbor a limited array of filamentous fungi and yeasts, following a succession of fungal species over time. Very little is known about the chemical ecology involved in the ensuing microbial interactions and in particular which secondary metabolites are produced and what role they play in developing the system. Such rotten or moldy apples are often removed from apple plantations, as well as from apple trees in private gardens. Of especial concern is the potential growth of *Monilinia* species, including species such as *M. fructigena, M. fructicola, M. laxa, M. yunnanensis,* and *M. polystroma*.(3, 4) Subsequent *Monilinia* conidium production can give rise to decay of further apples, if insects, birds, and other animals rupture the apple skin.(5) When these apples are left on the tree or on the soil they will give rise to the accumulation of *Monilinia* conidia. The development into black apples usually occurs after infection with several more fungal species after being left on the ground for some time. Even though black apples are not fit for human consumption, they constitute an interesting habitat where other microorganisms and animal species may live. In Nordic countries such as Denmark, apples infected with *Monilinia fructigena* will often also be infected with *Penicillium expansum* (blue mold rot), other species of filamentous fungi, and species of yeasts may also thrive in apples because of the facile availability of monosaccharides and disaccharides in apples.(6, 7) The low pH of apples, caused primarily by malic acid, will select for acidophilic filamentous fungi, which often produce specialized proteins, such as pectinases, glucose oxidase and hydrophobins,(7, 8) that play major roles in apple colonization. This may also result in the release of carbon sources that filamentous fungi can exploit for growth. For instance, *Monilinia* spp. produce pectinases (pectin lyases and polygalacturonases), proteases, xylanases, α-glucosidases and some species produce cutinases.(9) *P. expansum* and *P. crustosum* often grow on apples (with or without *Monilinia* growth), and *P. expansum* is well known for its production of the mycotoxin, quorum sensing inhibitor, and antibiotic, patulin.(10, 11) Several other species of *Penicillium* have been reported to grow on apples, including *P. crustosum* giving rise to a substantial rot, while species such as *P. solitum* and *P. polonicum* can cause minor rots in apples.(12–14) Apart from the rot on the surface of apples, other species of filamentous fungi can cause dry rot diseases in apples (at the core) including *Fusarium, Alternaria* and *Trichoderma* species.(6) Some of these fungi can inhibit the apple scab disease fungus *Venturia inequalis*.(15) However, these endophytic fungi do not produce a soft apple rot. Other species associated to rot of pomaceous fruits have included *Talaromyces minioluteus* and *T. rugulosus* (16) in addition to *Trichothecium roseum* (pink rot of apples).(17, 18)

Metataxonomic studies that have attempted to examine the fungal taxa of apple communities using next-generation high-throughput DNA sequencing to find both culturable and “non-culturable” species have proven to be inadequate for two main reasons.(19) Firstly, DNA sequence-based methods were used to detect the spora (all the spores present on the apple surface) rather than the actual apple funga (the fungi growing on apples).(19) Secondly, the fungi reported were mostly reported at the genus level, and thus neither *P. expansum* nor *Monilinia* spp. were reported from the apples examined by Shen *et al.*,(19) even though there is a vast literature on *P. expansum* and *Monilinia* spp. being the dominant soft rot fungi on the surface of apples. Furthermore, those that were listed as the “dominant” fungi at the species level have rarely, if at all, been reported as pathogens of apples. Whilst other microbiomes are harder to reproduce in the laboratory, this is less a problem for apples, and it should be feasible to isolate and cultivate all species naturally present.

*P. expansum* has been reliably reported to produce a large portfolio of secondary (specialized) metabolites: andrastin A-C, aurantioclavine, chaetoglobosins, citrinin (and related compounds such as 7-methyl-citrinin H1), communesin A-F, expansolide A-D, 5-farnesylisochromene, fumarylalanine, patulin, penexpandine, penostatins, piperafizin, roquefortine C, and N-formyl-roquefortine C,(11, 20–30) as well as the antibacterial specialized protein notatin,(31) and the necrosis-inducing protein NIP.(32) *Monilinia* spp. have been reported to produce moniliphenone, bromomonilicin, 4-bromopinselin, chloromonilicin, chloromonilinic acid A and B, chloropinselin, chrysophanol and emodin,(33–38) in addition to volatiles such as ethanol, dodecane and α-murolene.(39) Both *Penicillium* and *Monilinia* species also produce 1,8-dihydroxynaphthalene derived melanins.(40) *P. expansum* growing on apples produce the secondary metabolite patulin,(10) which has been reported to be an important factor for apple infection, but other secondary metabolites can also be important such as citrinin,(7, 41) and gluconic acid.(7) Acidification of the apple is needed to express sufficient endopolygalacturonase.(42) The role of fungal volatiles is less well-known, but *P. expansum* is known to produce geosmin and other volatiles,(43) some of which are even antifungal.(16)

In this study we investigated the chemical diversity and potential ecological role of microbially produced secondary metabolites in black apples – a unique and underexplored natural environment, one of high, yet restricted nutritional value, and of relatively low microbial diversity. We performed metabolomics analyses according to the protocol outlined in **Figure 1**. Using an OSMAC-guided approach, the fungal species present in black apples were determined chemotaxonomically, like the identity of individual compounds were assessed by analysis of HPLC-HRMS/MS data using spectral library matching tools such as GNPS, Dereplicator, as well as an in-house PCDL database. A focal point of this study involved how this relatively low microbial diversity was influenced by the dynamic production of microbial secondary metabolites, thereby shedding light on the role these metabolites might play for invasion and control of other microbes on black apples.

**Figure 1.**
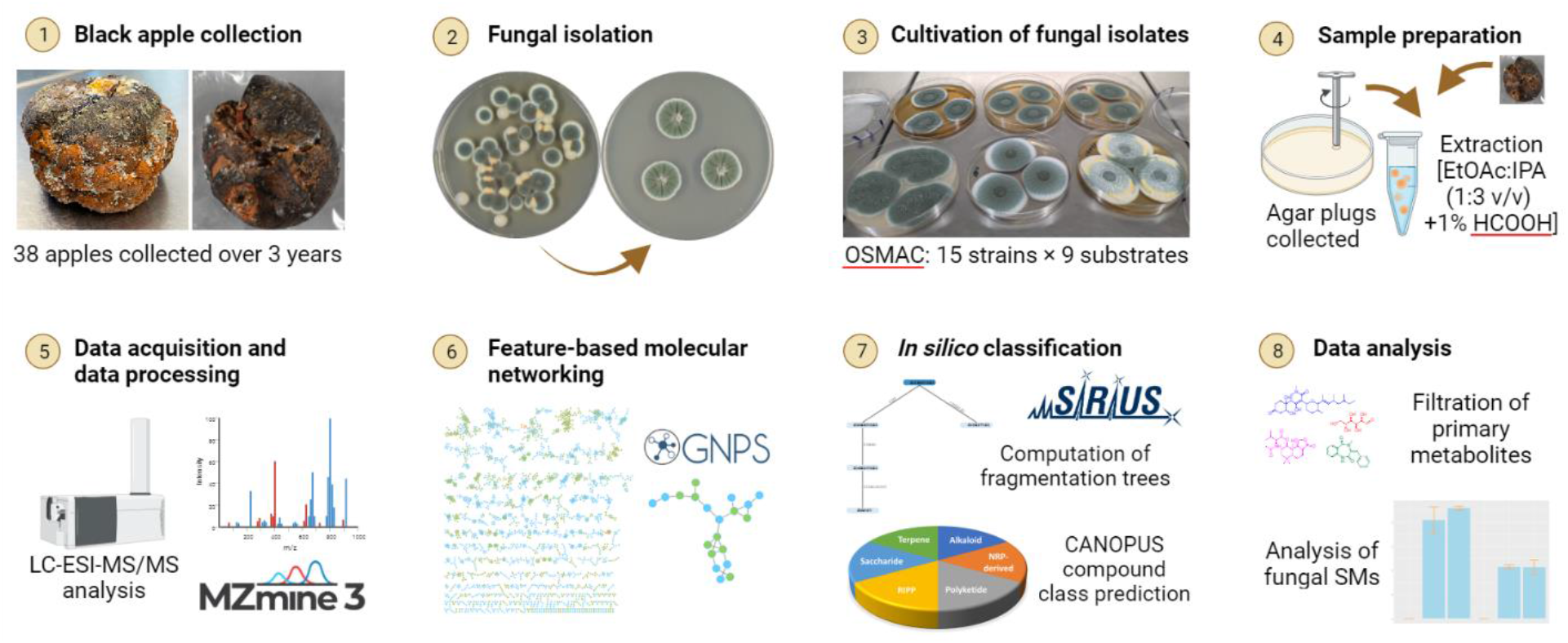
Workflow schematic for the metabolomics analysis of black apples. (1) Black apples were initially collected, (2) fungal strains were isolated, (3) fungal strains were cultivated on several media substrates, (4) agar plugs were collected for OSMAC study, and excisions collected of fungal biomass on black apples, followed by extraction using EtOAc:IPA (1:3 v/v) +1% HCOOH. Extraction was also conducted on whole black apples using the same solvent system. (5) The extracts were subjected to LC-ESI-HRMS/MS analysis in the positive ionization mode and processed in MZMine 3 for featuring finding and alignment of samples. (6) Feature-based molecular networking was conducted using GNPS, followed by (7) *in silico* classification of compound classes using the CANOPUS package in SIRIUS, with CSI-fingerID annotations computed also, and finally (8) all annotations merged into a unified table, compound class annotations propagated according to GNPS clusters, primary metabolites filtered out, and data analysis performed on fungal SMs identified in black apples.

## Results

### Key fungal species isolated from black apples and their associated secondary metabolites

The fundament of this study has been based on fungal cultures that were isolated from contaminated apples within a 30-year period (**Table S1**) and, in particular, fresh strains isolated recently from black apples. The list only contains relatively few *Monilinia* strains since our general research interest has been focused on the taxonomy and classification of species belonging to genus *Penicillium.* Chemical analysis was conducted on black apples collected during a three-year period (2019–2021), with samples A1-A29 corresponding to whole extractions of 29 different black apples collected in the first two years, and A30-A43 corresponding to small excisions (≈1 cm^2^) of fungal biomass collected across 9 apples in the following year, with in some cases, multiple species adjacent to each other on the same apple. From 6 of these more recent black apples, 10 fungal strains were isolated, corresponding to species of *M. fructigena*, *P. expansum*, *P. polonicum*, a *Penicillium* sp., *T. minioluteus* and *T. luteus*. These fungal isolates, supplemented with an additional 5 strains comprising *M. fructigena* and *P. expansum*, were profiled metabolically on an extended suite of media (nine substrates) in a one strain many compounds (OSMAC) approach (**Table S2**). The resulting metabolite profiles were used as an indicative pool of fungal secondary metabolites to query the whole black apples and apple-derived fungal excisions to determine which fungal secondary metabolites are important for invasion and control, and to chemotaxonomically determine the species present in black apples.

### Black apple associated fungi produce a huge diversity of secondary metabolites on artificial substrates

The OSMAC experiment revealed 3319 unique features associated with the 15 fungal isolates tested following the removal of features associated with blanks or primary metabolism. Of these features, 227 (6.8%) were attributable to known compounds as identified by a combination of GNPS, Depreplicator, CSI-FingerID, and PCDL results. From this pool of features, it was found that *P. expansum* represented the most talented producer of secondary metabolites, with 554 distinguishing features present in at least three samples across the four strains and nine media tested. It was determined that to minimize the potential for misidentification of features associated with a given species, a threshold presence of six indicative features consistent across at least 3 of the 27 samples for each isolate was required. There were also 370 distinguishing features identified in the *Penicillium* sp., 319 in *T. minioluteus*, 248 in *M. fructigena*, 156 in *P. polonicum*, and 153 in *T. luteus*.

In addition to the previously reported classes of chloromonilicinic acids (33, 35–37) (xanthones) and monilidiols (44) (partly-reduced monoaromatic oktaketides) from *Monilinia* spp., the production of a high-yielding, and uncharacterised, suite of secondary metabolites was observed (**Figures S1-S5**). These included the benzodiazepine 14-hydroxycyclopeptine ID: 9939, an unknown steroid as ID: 9126 (C_24_H_48_O_6_), and potential gluconic acid derived lipids (ID: 5675, C_18_H_36_O_6_; ID: 10766, C_26_H_52_O_6_; ID: 11025, C_25_H_50_O_5_). We also observed a new diketopiperazine from *P. expansum*, ID: 5762 (**Figures S6**). HR-ESI(+)-MS analysis revealed a protonated molecule indicative of a molecular formula C_18_H_18_N_2_O_2_ corresponding to the diketopiperazine, cyclo(Phe-Phe). MS/MS analysis revealed an immonium ion, *m*/*z* 120.0802, corresponding to the loss of phenylalanine. MS/MS fragmentation leading to the production of immonium ions are diagnostic of specific amino acid residues.(45) No other immonium ions or key product ions were detected, and with MS1 analysis indicating the presence of two phenylalanine residues, ID: 5762 was hence determined to be cyclo(Phe-Phe).

### A combined targeted and untargeted metabolomic approach highlights a high diversity of fungal secondary metabolites detectable in black apples

Following feature detection, dereplication and alignment of chemical features across the full dataset (both extracts of black apples and apple-associated fungal isolates), and the removal of features corresponding to the uncultivated media and primary metabolism, 4540 unique features were observed. Of the 1606 features observed in the black apples, 519 (32%) features could be directly assigned as being fungal-derived, due to their presence in the OSMAC-based metabolic profiling of the apple-associated fungal isolates. The remaining 1087 (68%) features observed in the black apples corresponded to natural apple metabolites, and fungal secondary metabolites only produced on apples or because of specific microbial interactions or degradative mechanisms. There were 90 (17%) features common to both the whole black apples and excisions, with 216 (42%) features unique to the whole black apples and 213 (41%) features unique to the excisions of black apples, indicating the complementary nature of the two datasets in providing a more exhaustive suite of fungal secondary metabolites in black apples. Of the features common to both the black apples and the apple-associated fungi, 54 (10%) features could be attributed to known compounds, with the first 27 (50%) assigned using the Global Natural Products Social (GNPS) Networking pipeline or through propagation of known classes within a GNPS cluster, a further nine (17%) being annotated using the Dereplicator tool on GNPS, a further five (9%) annotated using our in-house PCDL MS/MS fragmentation database, two (4%) were assigned based on comparison of MS/MS fragmentation reported in the literature, two (4%) were tentatively assigned based on MS/MS fragmentation predictions, and one (2%) predicted by CSI-FingerID (**Table 1**). Whilst the identification provided by MS/MS fragmentation matching is not as good as comparing to either the isolated compound by NMR, or the retention time of the standard under identical HPLC conditions, the tentative assignments ascribed here are of a very high level (B), according to the metabolite annotation guide proposed by Alseekh et al., 2021.(46) The remaining seven (13%) features were annotated to known compounds known to be produced by the producing species by HRESIMS level annotation and/or supplemented by investigation of isotopic patterns. This is the lowest level (D) of identification but combined with reported chemistry previously isolated in a given species is generally sufficient confirmation. Whilst patulin was not detected in any of the naturally infected black apples computationally, the biosynthetic precursor, and its isomer, (*E*)- and *(Z*)-ascladiol were present.(47) These derivatives are also known biotransformation products of patulin by certain yeasts and bacteria.(48, 49)

**Table 1.**
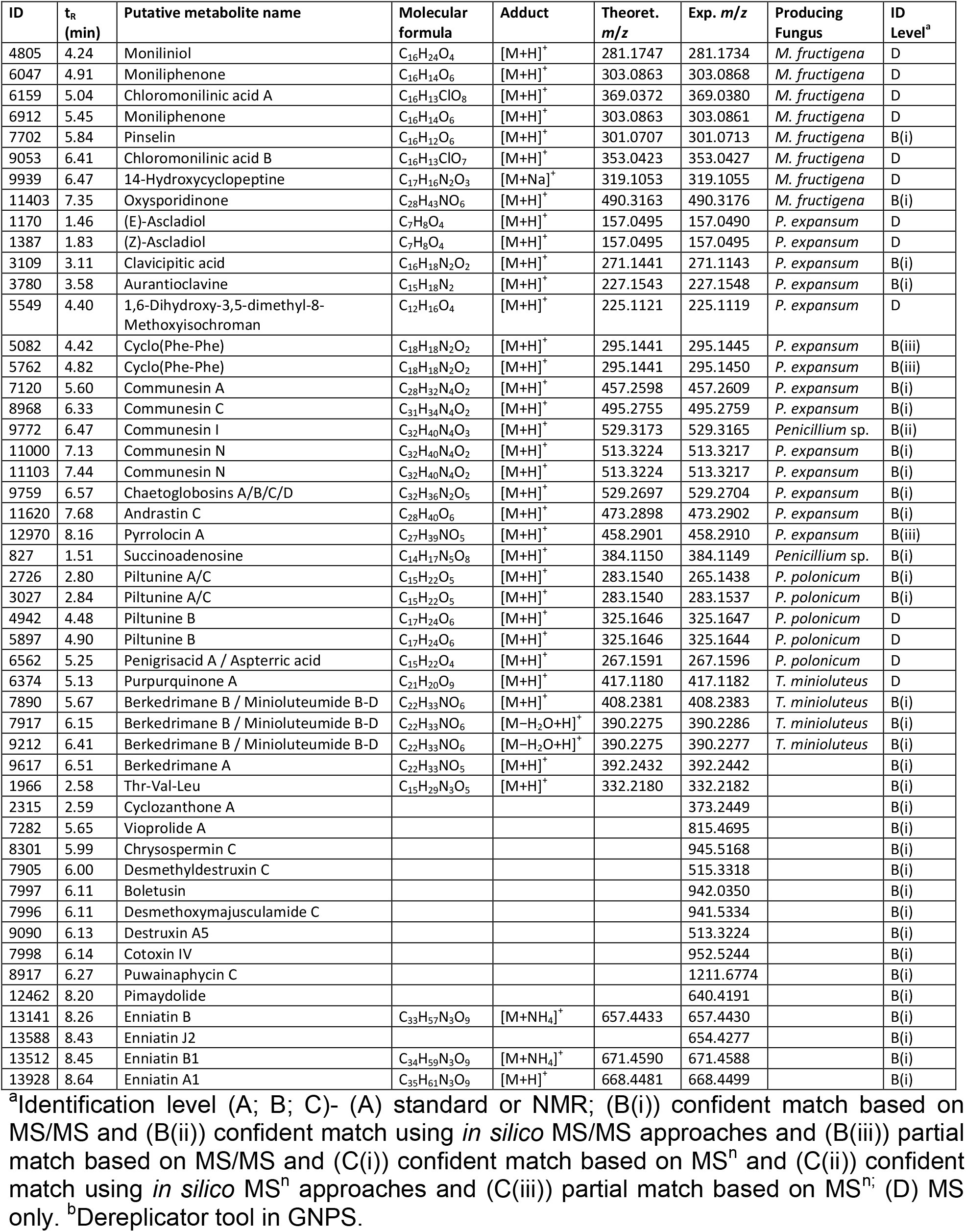
Annotations of screened fungal secondary metabolites identified in black apples.

### *Monilinia fructigena* produces mainly NRPS derived compounds on black apples

Due to the vast pool of unknown secondary metabolites detected, categorisation of the specific classes of compounds being produced on black apples was computed, first by using SIRIUS (50) to predict formulae with the ZODIAC tool (51) followed by prediction of the compound class with the CANOPUS tool (52–54) based on comparison of MS/MS fragmentation patterns (**Figure 2**). Surprisingly, *M. fructigena* was shown to produce mostly NRPS-derived compounds, with small peptides accounting for 43% of the features, followed by oligopeptides (22%) rather than the often reported two major types of polyketides. The suite of identified features identified in black apples from *M. fructigena* included the described diketopiperazines, gluconic acid derived lipids and a high abundance of the unknown steroid, ID: 9216. Compound classes for the other two genera, *Penicillium* and *Talaromyces*, were more diverse including sesquiterpenoids (e.g. berkedrimane B), tryptophan alkaloids (e.g. communesins) and cyclic polyketides (e.g. ascladiols).

**Figure 2.**
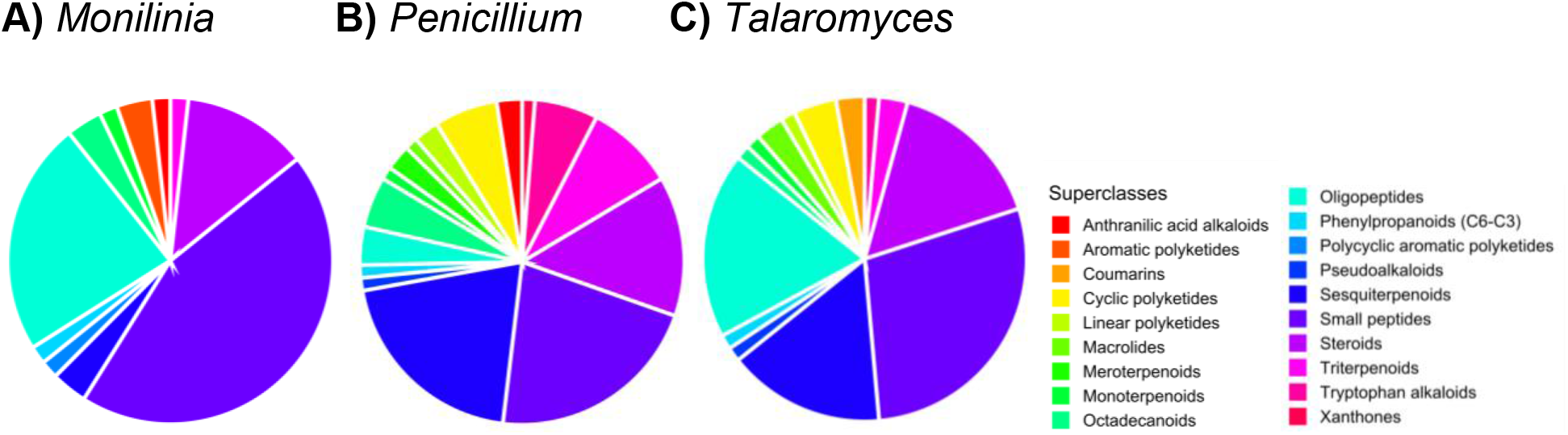
Proportion of fungal secondary metabolites found in black apples associated with the relevant natural product superclasses (>70% confidence predicted by CANOPUS tool in SIRIUS) for the genera: A) *Monilinia*, B) *Penicillium*, and C) *Talaromyces*.

In addition to comparing the differences in compound classes produced by the different fungal genera on black apples, the differences in compound classes between the different black apple samples was also compared (**Figure 3**). Notably, the presence of certain compound classes differed between the whole black apples (A1-A29) and the excisions of black apples (A30-A43), with cyclic and polycyclic polyketides (such as ascladiol and chloromonilinic acid B produced by *P. expansum* and *M. fructigena*, respectively) only detectable in the former, and anthranilic acid alkaloids only detectable in the latter. The differences noted here could be due to the localised concentration differences between secondary metabolites or the presence of potentially different fungal species based on the different locations from which the black apple samples were derived.

**Figure 3.**
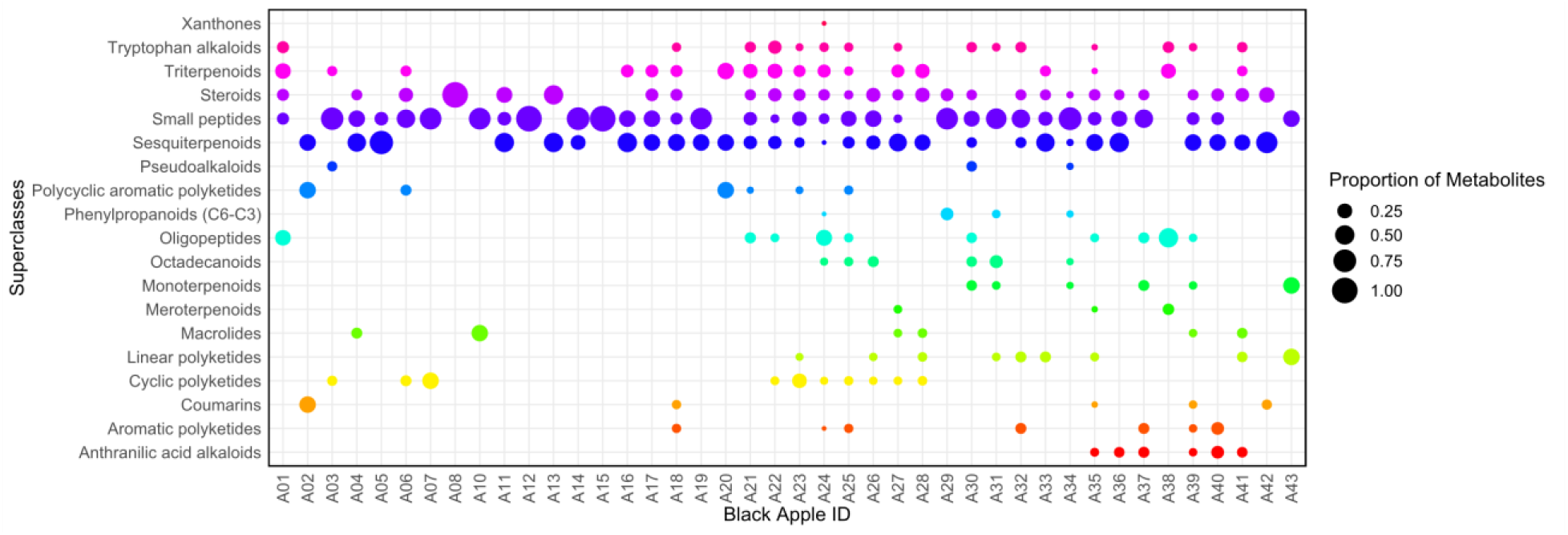
Proportion of fungal secondary metabolites associated with the relevant natural product superclasses for each black apple, shedding light on secondary metabolite classes of ecological significance in invasion and/or control of apples.

### Relative abundances of fungal secondary metabolites indicate those ecologically important for invasion and control of black apples

The ecological importance of certain secondary metabolites to the black apple system was ascertained by comparing the average normalised abundances produced on black apples to those produced in the laboratory on standard cultivation media. Normalisation was performed for each sample according to the maximum chromatographic peak area. Interestingly, whilst it was found that known secondary metabolites tended to be produced at equivalent levels or was suppressed in the black apples compared to that observed by the OSMAC approach (Table S4), there were some notable exceptions. Namely, the antifungal compound, berkedrimane B,(55) produced by *T. minioluteus* was found to be 28 times higher in black apples on average compared to on standard cultivation media. Pinselin and moniliphenone, biosynthetic precursors to the antifungal compound, chloromonilicin,(37) by *M. fructigena* were also 15-25 times more abundant in black apples. Whilst the *Monilinia*-derived chloromonilicin and chloromonilinic acid A were not detected in the OSMAC study as per the LCMS analysis conducted in the positive ionization mode, the compounds were identified in 9 and 20 of the black apple extracts, respectively. This indicates that these antifungal compounds from *T. minioluteus* and *M. fructigena* are important for competitive invasion and control of other microbes on black apples.

For *P. expansum*, it was found that whilst andrastins A/B, citrinin, expansols A/B, expansolides A/B, and roquefortine C, are highly produced in the OSMAC study, they were not observed in the black apples. This perhaps highlights compounds which are not important in this ecological niche or are rapidly biotransformed or degraded. This could be due to the presence of these compounds being more abundant in spore-containing regions of *P. expansum*, which are often no longer present on older (mummified) black apples.

Fungal excisions were also compared as a means of investigating the potential microbial interactions in a defined spatial resolution of a given black apple, with for example the adjacent A32 (excision of *M. fructigena*) and A33 (excision of *T. minioluetus*) highlighted in **Figure S7**. In this case, it was found that the major differences in the abundance of produced secondary metabolites between excisions was mainly observed for uncharacterised metabolites. It is tempting to speculate that such unknown compounds are related to biosynthetic gene clusters, only expressed in the natural system, however more investigations are needed to prove this hypothesis.

### Metabolic profiling of black apples distinguishes the presence and temporal succession of fungi

Of the 43 black apple extracts obtained in this study, there were 30 instances where fungal secondary metabolites belonging to one of the key species investigated by the OSMAC approach could be identified (**Figure 4**), based on the scoring system applied here. Unsurprisingly, metabolites pertaining to *M. fructigena* signified the bulk of the black apples, with 26/30 indicating its presence. This gives credence to the speculated succession order for the invasion of fungal species giving rise to black apples, with *M. fructigena* generally observed as the initial coloniser of apples, indicated by brown ringlets on their surface. The next most commonly observed fungal species based on their chemical “footprints” were *P. expansum* and *T. minioluteus*, accounting for 21 and 20 of the black apples extracted, respectively. To find the presence of secondary metabolites belonging to *T. minioluteus* in such a high proportion of the black apples was indeed surprising, given its reported isolation is much lower. This is followed by the *Penicillium* sp., *P. polonicum* and *T. luteus.* The remaining 13 black apples where no determinable fungal species could be identified are quite peculiar, with possible biotransformations of known chemistry, inductions of novel chemistry from known competitors, or the potential presence of more obscure and possibly uncultivable species. However, it is noted that it was evident that the two known metabolites, pyrrolocin A and berkedrimane B, produced in our queried species were identified but collectively with other metabolites did not meet our threshold of six indicative metabolites needed to be present for confirmed presence of a species in each black apple. Choosing a lower threshold of three indicative features, for instance, altered the detected frequency for metabolites belonging to *M. fructigena*, where only three samples indicated no prior growth (data not shown).

**Figure 4.**
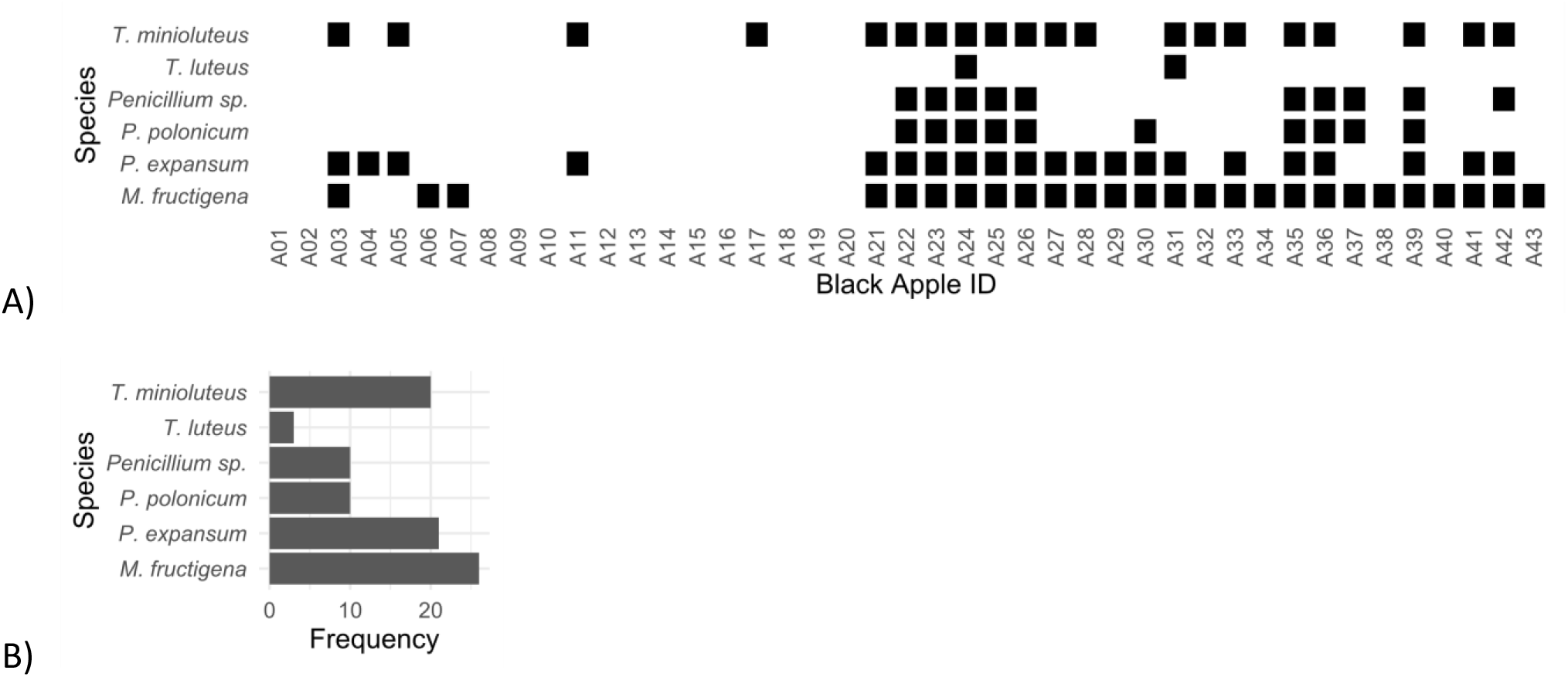
A) Heatmap showing the presence of each fungal species in each black apple; and B) barchart showing the associated frequency of each species present in black apples.

Interestingly, there was a single black apple, A24, that showed evidence for secondary metabolites belonging to all six of the fungal species investigated by the OSMAC approach. LCMS/MS analysis of this apple indicated it to have the highest degree of chemical complexity, with the chemical extract containing 311 features pertaining to secondary metabolites, approximately 3-fold higher than the mean (109 ± 63), and indeed, far higher than that pertaining to the mean of the black apples where no OSMAC-derived fungal secondary metabolites were detected (81 ± 40). The major metabolite present is a likely flavonoid with the predicted formula, C_15_H_10_O_8_, which we highlight here due to its absence in any of the blank media or natural apple substrates, and hence could be a biotransformation product. Outside of black apples, the compound was most observed for our single strain of *T. luteus* on rice substrates.

Outside of the 15 strains selected for OSMAC analysis were other secondary metabolites from other fungal species detected in black apples, namely: mycophenolic acid, a highly biologically-active metabolite found *Penicillium brevicompactum* and other fungi,(56–58) aurofusarin, from the wheat pathogen *Fusarium* sp.,(59), enniatins from *F. culmorum*, the structurally-related xanthoepocin, a common metabolite from the *Penicillium* genus,(60, 61) asperphenamate, derived from several Aspergillaceae,(62–66) and berkedrimane A originally described from a *P. solitum* strain.(67) However, we speculate that the latter strain was misidentified, since we have never found berkedrimanes to be produced by any *P. solitum* strain.(68) Instead, we have consistently detected berkedrimanes from the *Talaromyces* species *T. amestolkiae*, *T. purpureogenus* and *T. trachyspermus*.(69) Interestingly, the presence of mycophenolic acid, xanthoepocin and asperphenamate together, are key indicators that *P. brevicompactum* may have been present on six of the apples (A3, A6, A7, A23, A25, and A27). Notably, all six of these apple extracts are from whole black apples. Furthermore, the presence of aurofusarin, like in A24 and A38, as well as enniatins only in A24, could indicate chemical footprints of the dry rot fungal plant pathogens, *Fusarium* sp., species often also commonly associated with the plant leaves or stems.

### Networking using GNPS identifies ecologically important secondary metabolite inductions, decompositions and biotransformations in black apples

For *P. expansum*, the communesin class of secondary metabolites, which have reported modest cytotoxic activity,(25, 70) demonstrated considerable chemical diversity across the entire dataset with the presence of communesins A, C, I, and N, the previously identified, but not characterised, Com622,(26) and two unknown communesins (**Figure 5a**). Rather than the communesin analogues being produced together, each communesin analogue seemed to be specifically produced in unique black apples, or along with only one other analogue, suggesting a microbiome-dependent production by *P. expansum*. For instance, communesin A and the unknown communesin with *m*/*z* 538 were found only in the black apple chemical extract A30, communesin C only in A32, and Com510 only in A22. Surprisingly, it was the tentatively identified and only very recently characterised analogue, communesin N,(25) with the highest frequency among black apples, with presence in four black apples (A1, A21, A22, and A24). Communesin N is most structurally similar to communesin B, with reduction of its amidyl side-chain, and hence its presence could be due to the high abundance of antioxidants in apples. There was also an unknown communesin with *m*/*z* 511 that could correspond to compound 7a in the paper by Hoang, *et al*., a partially reduced intermediate between communesin B and N.(25) In general, it is likely that the chemical diversity of communesins observed in black apples arises from upregulation of certain analogues by *P. expansum*, or chemical degradation by pH and apple flavonoids.

**Figure 5.**
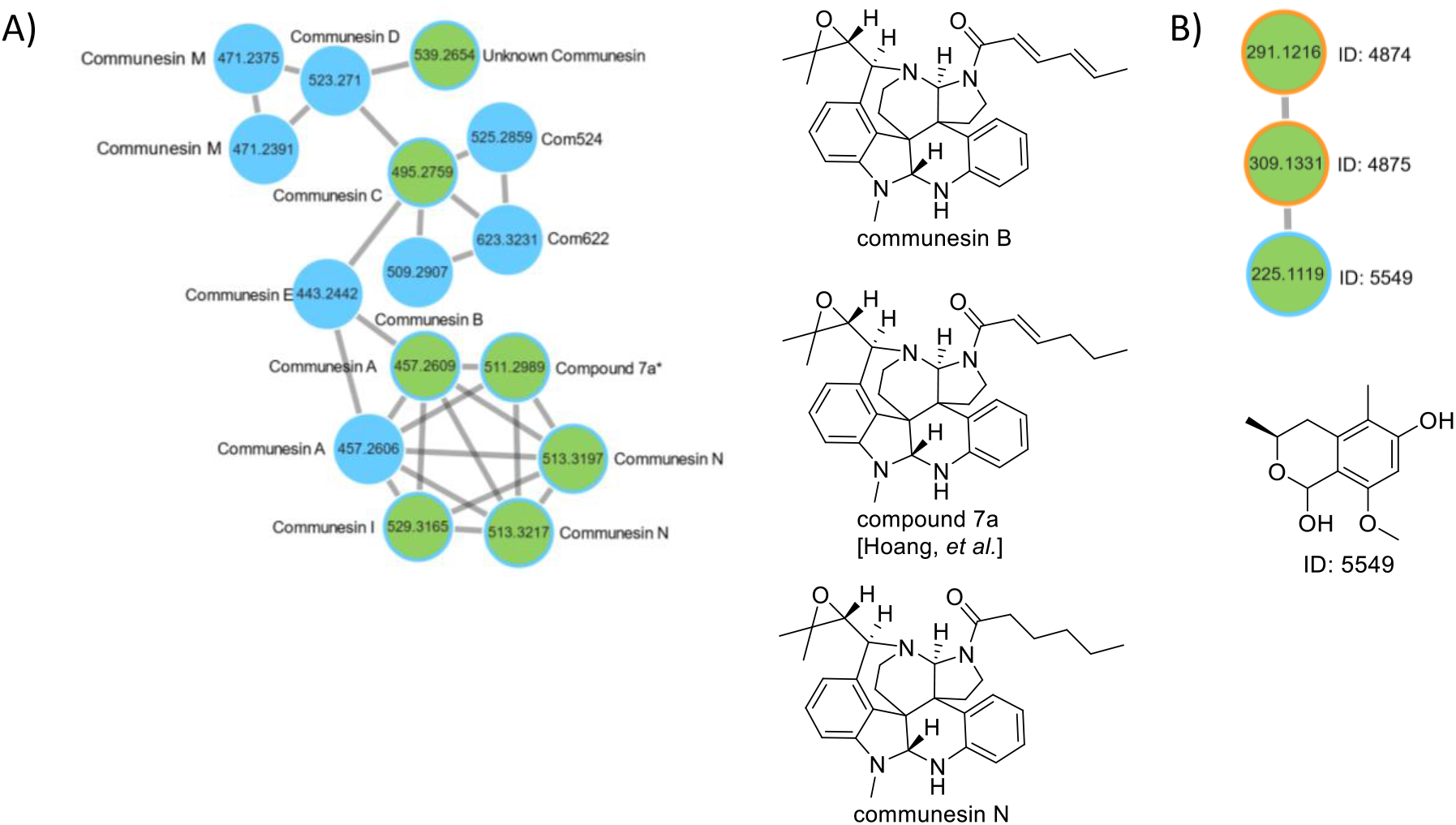
Molecular family of metabolites belonging to the class of A) communesins and B) isochromans derived from the black apple network (GNPS). Nodes corresponding to fungal features identified (presence/absence) in black apples are filled in green, and those absent in blue. The blue border Additionally, nodes that are identified in the OSMAC study of the 15 selected fungal isolates have a blue border, and those that are unique present in the black apples with an orange border. ID: 4874 and ID: 4875 have the same retention time and hence correspond to the same compound.

Networking using GNPS indicated the biotransformation, induction, or degradation of several other fungal secondary metabolites. For instance, the identified isochromenone ID: 2983, C_10_H_10_O_5_ produced by the *Penicillium* sp. on jasmine rice, shared a cluster with a hydrolysed analogue ID: 3699, C_10_H_8_O_4_, which was only observed in the black apples. Further hydrolyses can be seen for GNPS clusters 247 and 484. Reductions of fungal secondary metabolites were also very common, with conversion of ID: 2015, C_15_H_18_O_3_ produced by *P. expansum* only on basmati rice to ID: 5363, C_15_H_20_O_3_ in the black apples and the reduction of ID: 14961, C_34_H_56_O_9_ produced by *T. minioluteus* also only on basmati rice to ID: 15389, C_34_H_58_O_9_ in the black apples. One potential induction observed in the black apples, as determined by clustering in GNPS, involved the appearance of the isochroman, ID: 4875, C_16_H_20_O_6_, related to the known ID: 5549 (1,6-dihydroxy-3,5-dimethyl-8-methoxyisochroman) produced by *P. expansum* (**Figure 5b**). Whether the production of this new isochroman – a hypothetical esterified product at its secondary hydroxy group – was due to the presence of other microbial partners, the specific apple substrate, or a biotransformation product is yet to be determined.

Whilst the aim here was to focus more on the microbial secondary metabolism associated with black apples, we would be remiss to ignore the secondary metabolism of apples in general. Common apple secondary metabolites which were observed included (+)-catechin, (−)-epicatechin, quercetin, kaempferol, ursolic acid, chlorogenic acid, and other flavanols, phenolic acids, terpenes, carotenoids and organic acids.(71) No obvious microbial biotransformations of known apple secondary metabolites were detected.

## Discussion

In this study, the microbial secondary metabolite chemical diversity in black apples was investigated by linking features present in apple-associated fungal isolates grown on standard cultivation media to those detectable in black apples. In so doing, we were able to chemotaxonomically predict which species might have been present in a given black apple and link the produced secondary metabolites to potential ecological roles in invasion or control of a given black apple. Here, it was observed that at least one of our six species selected for OSMAC experiments was present in 30/43 of the black apple extracts examined, based on the criteria of detection of at least six representative chemical features of our key species. The likely presence of additional untested fungi, such as *Fusarium* sp. and *P. brevicompactum*, can also be seen as determined by the presence of indicative metabolites. Although *M. fructigena* and *P. expansum* were confirmed to be the two most prolific fungal species present on black apples, the frequency of detection of metabolites associated with *T. minioluteus* (20/43 of the black apple extracts) was higher than expected based on that observed for the literature. This could be due to the low number of reports associated with black apples, or it could be a phenomenon more geographically linked to apples produced in Denmark (as indicated by the frequency of isolation in Table S1). Thus, we have shown that the individual microbial and chemical diversity originating from black apples to be underrepresented, indicating a complex network of chemical cues important for establishment, control, and homeostasis. Because of this relatively high observed chemodiversity, the black apple model system represents an interesting environment for the study of the ecological role of secondary metabolites.

Whilst the exact ecological role of specific secondary metabolites is more difficult to discern from such a generalised study, it can be seen that many of the known metabolites – which also tend to correspond with those that are highly represented on standard cultivation media – are present in black apples at significantly lower levels than expected, highlighting the specificality of the apple substrate and the interactions between the indicated microbial partners to be highly influential to the expressed apple microbiome secondary metabolome. However, the antifungal or antifungal precursor compounds, berkedrimane B, pinselin and moniliphenone, are present in much higher relative abundances in black apples, highlighting them as potentially important compounds for invasion and control of black apples by the producing fungi. In general, our results indicate that production of secondary metabolites on black apples is tightly regulated, influenced by certain environmental cues (such as low pH), stimuli, and limitations due to the added fight for resources with other competitors present.

One explanation for the suppressed production of certain metabolites is that they are being detoxified. Antioxidants, for instance, are known to assist in the detoxification of potentially harmful molecules,(72) and are highly abundant in the form of plant flavonoids. In fact, this is perhaps reflective of a dual role for plant flavonoids, such as catechin found in apples, for both antibiosis and detoxification of fungal secondary metabolites.(73) Previous studies have shown that apples high in procyanidins, dihydrochalocone, flavonols, and hydroxycinnamic acids were more resistent to infection by *P. expansum*.(74) When mycotoxin producers are treated with phytochemicals, the production of mycotoxins is significantly reduced. In *Aspergillus flavus*, for instance, aflatoxin production has been shown to be decreased by up to 99.8% with the addition of hydrolysable tannins, flavonoids and phenolic acids derived from pine nuts.(75) Hence, we hypothesise that the presence of natural antioxidants in apples is likely an influential reason for the lower than expected metabolic output for known fungal secondary metabolites in black apples.

Not only are fungal secondary metabolites influenced by plant phytochemicals, but also the secondary metabolites of other competing species, which may oxidise and hence nullify the chemical repertoire of a fungus. Furthermore, the chemical fate of secondary metabolites can be even more complex, with the low pH of apples (often around pH 3) causing chemical degradation and biotransformations between interacting species. Such biotransformations have also been linked to detoxification mechanisms, and most commonly observed for antibiotic resistance(76) and quorum quenching (77–79) mechanisms. Some possible biotransformations of fungal secondary metabolites in black apples have been shown here, with hydrolysis or reduction most observed, although there is the possibility this is due to the low pH environment of apples. To investigate the mechanism behind these chemical transformations, one could take the purified known compounds and systematically treat the compounds with relevant fungal species, or acidified solutions, and with the aid of GNPS clustering determine if the conversion is enzymatic or non-enzymatic. A recent study by Kang, K. B and coworkers systematically investigated the potential biotransformations of phytochemicals by a panel of fungi using GNPS.(80)

Another surprising observation was the high abundance of compounds associated with *M. fructigena* present, including the uncharacterised steroid, ID: 9216, as well as several NRPS derived diketopiperazines and gluconic acid derived lipids. Gluconic acid has previously been shown to have a role in the virulence of the related *M. fructicola* on peaches,(81) and also by *P. expansum* on apples,(82) and is a means of locally modifying the pH of their environment. On the other hand, the presence of diketopiperazines perhaps reflects on their antimicrobial and quorum sensing inhibition roles,(83) which is notable given these metabolites are also highly abundant in black apples. Further investigations should investigate both the structural diversity and the potential ecological role these compounds might play in the invasion and control of *M. fructigena* on black apples.

In conclusion, this metabolomics study of black apples has provided many new insights into the presence and potential ecological roles of microbially, and in particular fungal, produced secondary metabolites in a natural system. Such a microbiome is highly advantageous for the study of secondary metabolites due to the high chemical diversity detectable from a relatively small selection of acidophilic microbial species, and the largely reproducible succession of microbes that is observed annually. The interactions between the invading microbes are dynamic and the resulting black apple metabolome a reflection of the key secondary metabolites which are important for invasion, control and temporal fungal progression within such a system. Whilst many of the known metabolites were shown to be diminutively observed, perhaps due to detoxification and degradative mechanisms of phytochemicals and other highly reduced fungal secondary metabolites, enzymatic processes in interacting species, and the low pH environment, there was also a high repertoire of upregulated metabolites arising from inductions and biotransformations. To understand this system further, it will be important to investigate the interactions on a more simplified level, delving deeper into the biological and ecological role of these more highly abundant molecules that are produced in dual cultures(84) and synthetic communities (SynComs) as has been recently created for many other microbiomes.(85–87) A potential SynCom that would be important to investigate could include a combination of yeasts and the filamentous fungi, *M. fructigena*, *P. expansum* and *T. minioluteus*.

## Methods and Materials

*Isolation of apple-associated fungi.* Fungal strains were isolated either by classical collection and transfer of fungal spores from black apples into a defined spore solution, followed by streaking (and potentially re-streaking) onto agar plates, and cultivation at room temperature for 5-7 days, until a pure culture was obtained. Alternatively, fungal cultures were isolated by crushing of mummified apples, followed by spreading on V8 and CYA with antibiotics. Spore suspensions were made by harvesting spores using 20% glycerol, and subsequently storing them in −80C. All cultures are kept in the IBT collection (https://www.bioengineering.dtu.dk/research/strain-collections/ibt-culture-collection-of-fungi).

### Cultivation for metabolic profiling of fungal isolates

Metabolite profiling was conducted by inoculating fungal isolates onto yeast extract and sucrose (YES), Czapek yeast extract (CYA), malt extract (MEA – DIFCO and OXOID), oatmeal (OAT) agar, potato dextrose agar (PDA), jasmine rice, basmati rice, and cracked wheat. *M. fructigena* was cultivated at 20 °C under alternating light conditions (12 h light, 12 h dark) for 7, 14 and 21 days. The remaining fungi were cultivated at 25 °C in darkness for 7, 14 and 21 days.

### LC-MS/MS sample extraction and preparation

All samples used for extraction were HPLC grade or better. For fungi cultivated on agar, 3 plugs (≈8 mm in diameter) were added to an Eppendorf tube prior to extraction. For fungi cultivated on grains, grains and mycelia totalling a volume of ≈0.5 cm^3^ were added to an Eppendorf tube prior to extraction. For excisions of fungi on black apples, excisions of the fungus and shallow penetration of the apple exterior totalling a volume of ≈0.5 cm^3^ were added to an Eppendorf tube prior to extraction. In each of these cases, the samples were extracted in isopropanol/ethyl acetate (1:3) + 1% HOOH (1 mL). For extractions of whole black apples, the apples were pureed using an IKA dispenser. If needed, a small volume of Milli-Q water was added during homogenization. Following homogenization, the metabolites were extracted with equal volumes (≈3 mL) of isopropanol/ethyl acetate (1:3) with 1% formic acid in a 15 mL falcon tube. The resulting chemical extracts were sonicated for 30 min, and the supernatants dried under nitrogen. Dried samples were resuspended in methanol (1 mL) and centrifuged before analysis by UHPLC-DAD-MS.

### Data-Dependent LC-ESI-HRMS/MS Analysis

Ultra-high-performance liquid chromatography– diode array detection–quadrupole time-of-flight mass spectrometry (UHPLC–DAD–QTOFMS) was performed on an Agilent Infinity 1290 UHPLC system (Agilent Technologies, Santa Clara, CA, USA) equipped with a diode array detector. Separation was achieved on a 150 × 2.1 mm i.d., 1.9 μm, Poroshell 120 Phenyl Hexyl column (Agilent Technologies, Santa Clara, CA) held at 40°C. The sample (1 μL) was eluted at a flow rate of 0.35 mL min^−1^ using a linear gradient from 10% acetonitrile (LC-MS grade) in Milli-Q water buffered with 20 mM formic acid increasing to 100% in 10 min, staying there for 2 min before returning to 10% in 0.1 min. Starting conditions were held for 3 min before the following run.

Mass spectrometry (MS) detection was performed on an Agilent 6545 QTOF MS equipped with Agilent Dual Jet Stream electrospray ion source (ESI) with a drying gas temperature of 250°C, a gas flow of 8 L min^−1^, sheath gas temperature of 300°C and flow of 12 L min^−1^ Capillary voltage was set to 4000 V and nozzle voltage to 500 V in positive mode. MS spectra were recorded as centroid data, at an *m/z* of 100–1700, and auto MS/HRMS fragmentation was performed at three collision energies (10, 20, and 40 eV), on the three most intense precursor peaks per cycle. The acquisition rate was 10 spectra s^−1^. Data were handled using Agilent MassHunter Qualitative Analysis software (Agilent Technologies, Santa Clara, CA). Lock mass solution in 70 % MeOH in water was infused in the second sprayer using an extra LC pump at a flow of 15 μL min^−1^ using a 1:100 splitter. The solution contained 1 μM tributylamine (Sigma-Aldrich) and 10 μM Hexakis (2, 2, 3, 3-tetrafluoropropoxy) phosphazene (Apollo Scientific Ltd., Cheshire, UK) as lock masses. The [M + H]^+^ ions (*m/z* 186.2216 and 922.0098, respectively) of both compounds were used.

### LC-MS/MS data processing

The mass spectrometry data were centroided and converted from the proprietary format (.D) to the *m*/*z* extensible markup language format (.mzML) using ProteoWizard (version 3.0.22112, MSConvert tool).(88) The mzML files were then processed with MZmine 3 toolbox.(89) The .mzML files, metadata (.txt), MZmine batch file (.XML format) and results file (.MGF) are available in the MassIVE dataset MSV000092823. The MZmine batch file contains all the parameters used during the processing. Namely, feature detection and deconvolution was performed with the ADAP chromatograph builder (90) and local minimum search algorithm. Isotopologues were then identified and filtered, and the remaining features aligned across samples. To reduce the number of features resulting from the same molecule, duplication filtering, metacorrelate and ion identity networking was performed. For the final feature quantification table, a.MGF file was exported in both GNPS (91) and SIRIUS relevant formats as well as a .CSV file. We also provide complete computational workflows for the metabolomics analyses conducted (available on GitHub at https://github.com/michael-cowled/black-apples).

### LC-MS/MS data annotation

In order to dereplicate known compounds in the chemical extracts, the raw MS/MS spectra were searched against the in-house library using Agilent MassHunter PCDL manager (Agilent Technologies) (92). Due to the manual nature of this task, this was only carried out for a randomly selected 10% of the samples.

The remaining annotations were performed using the processed mass spectrometry data (.MGF and .CSV files from MZmine). Firstly, GNPS was performed using the feature-based molecular networking workflow. (91) In addition, the DEREPLICATOR tool was used for the annotation of small peptides. SIRIUS (v 5.7.0) (50) was used to compute molecular formulae, through comparison of experimental and predicted isotope patterns,(93) computation of fragmentation trees,(94) and further improved with the ZODIAC module.(51) CSI:FingerID (95, 96) was employed to conduct *in silico* structure annotation utilising structures from biodatabase, which allowed for compound class categorisation with CANOPUS.(52–54) SIRIUS was configured with a mass accuracy of 10 ppm and for annotation of the ions [M+H]^+^, [M+K]^+^, and [M+Na]^+^. In the case of ZODIAC, the features were divided into 10 random subsets using GIBBS sampling to alleviate computational burden. Each subset was computed separately, employing a threshold filter of 0.95 and requiring a minimum of 10 local connections.

In order to identify potential microbially produced secondary metabolites, we gathered annotations from spectral library matching (in GNPS) and the DEREPLICATOR+ tools, as well as through querying our in-house PCDL database.(92) These annotations were subsequently compared against the Natural Products Atlas v2021_08 (97) and MIBiG v3.1 (98) databases pertaining to secondary metabolites. Compound class annotation for unknown compounds was appended from NPClassifier for which the NPC superclass probability ≥ 70%. Filtration of primary metabolite classes was carried out on the resulting annotated dataset. The workflow is akin to, and hence was inspire by, that used in a recent report for the Earth Microbiome Project 500 (EMP500), except for propagation of compound classes using GNPS clusters, as we found this removed several features of which we deemed important.(99)

### LC-MS/MS data analysis

Following combination of the above annotations, filtering and computations, the resulting table of features was linked with the corresponding metadata associated with the samples. Features present in blank media controls were subtracted from the dataset if present above a threshold of 1000. At this point, the data was split in accordance with whether the samples were derived from black apples, or from fungal cultivation of standard laboratory media (OSMAC). The resulting two datasets were compared, focussing on the features present in both. Each apple extract was also queried using the suite of metabolites identified in each species (present in at least 3 samples associated with the species by OSMAC) in order to assign its presence in a given black apples sample with at least 6 features associated with a species needed in order to confirm its presence. All plots were generated in R using the ggplot2 package.(100)

## Acknowledgements

This study was carried out as part of the Center of Excellence for Microbial Secondary Metabolites (CeMiSt) funded by The Danish National Research Foundation (DNRF137). We thank the members of CeMiSt for general scientific discussions and advice.

